# Triku: a feature selection method based on nearest neighbors for single-cell data

**DOI:** 10.1101/2021.02.12.430764

**Authors:** Alex M. Ascensión, Olga Ibañez-Solé, Inaki Inza, Ander Izeta, Marcos J. Araúzo-Bravo

**Affiliations:** Biodonostia Health Research Institute, Computational Biology and Systems Biomedicine Group, Paseo Dr. Begiristain, s/n, 20014, Donostia-San Sebastian, Spain; Tissue Engineering Group, Biodonostia Health Research Institute, Paseo Dr. Begiristain, s/n, 20014, Donostia-San Sebastian, Spain; intelligent Systems Group, Computer Science Faculty, University of the Basque Country, Donostia-San Sebastian, Spain; Max Planck Institute for Molecular Biomedicine, Roentgenstr. 20, 48149, Muenster, Germany

**Keywords:** scRNAseq, feature selection, bioinformatics, python

## Abstract

Feature selection is a relevant step in the analysis of single-cell RNA sequencing datasets. Triku is a feature selection method that favours genes defining the main cell populations. It does so by selecting genes expressed by groups of cells that are close in the nearest neighbor graph. Triku efficiently recovers cell populations present in artificial and biological benchmarking datasets, based on mutual information and silhouette coefficient measurements. Additionally, gene sets selected by triku are more likely to be related to relevant Gene Ontology terms, and contain fewer ribosomal and mitochondrial genes. Triku is available at https://gitlab.com/alexmascension/triku.

## 1 Background

Single-cell RNA sequencing (scRNA-seq) is a powerful technology to study the biological heterogeneity of tissues at the individual cell level, allowing the characterization of new cell populations and cell states–i.e. cell types responding to different environmental stimuli– previously undetected due to their low frequency within the tissue and the lack of individual resolution of bulk methods [1, 2].

scRNA-seq datasets are multidimensional, i.e. the expression profile per cell consists of multiple genes. Two common characteristics of multidimensional datasets is their high dimensionality and their sparsity, which are worsened in single-cell datasets due the high proportion of zeros from low signal recovery [3]. This sparsity affects downstream methods such as cell type detection or differential gene expression [4].

A common task when working with multidimensional datasets is feature selection (FS). FS, alongside with feature extraction (FE), responds to the need of obtaining a reduced dataset with a smaller dimensionality [5]. While FE methods like Principal Component Analysis (PCA) extract new features based on combinations of the original features, FS methods aim to select a subset of the features that best explains the original dataset.

There are three main types of FS methods: filter, wrapper and embedded methods [5]. Current FS methods in scRNA-seq analysis are filter methods because common downstream analysis steps do not embed the FS within the pipeline [6]. FS methods represent a key step in processing pipelines of bioinformatic datasets and provide several advantages [5]: they reduce model overfitting risk, improve clustering quality, and favour a deeper insight into the underlying processes that generated the data (features –genes– that contain random noise do not contribute to the biology of the dataset and are removed). Specifically, in scRNA-seq, removing non-informative features can improve results in downstream analyses such as differential gene expression.

Early methods for FS in scRNA-seq data were based on the idea that genes whose expression show a greater dispersion across the dataset are the ones that best capture the biological structure of the dataset. Conversely, genes that are evenly expressed across cells are unlikely to define cell types or cell functions in a heterogeneous dataset. The most straightforward way of selecting genes that are not evenly expressed is to look at a measure of dispersion of the counts of each gene and to select those genes that have a dispersion over a threshold.

However, the correlation between mean expression and dispersion introduces a bias whereby genes with higher expression are more likely to be selected by FS methods. However, biological gene markers that define minor cell types are usually expressed in a medium to small subset of cells. Therefore, new FS methods based on dispersion are designed to correct for this dispersion/expression correlation to select genes with a broader expression spectrum.

Brennecke et al. [7] developed a FS method that introduces a correction over the dispersion that accounts for differences in the mean expression of genes. It does so by setting a threshold to the correlation between the average gene expression and its coefficient of variation across cells. Newer FS methods have arisen after different corrections, like the one originally described by Satija et al. [8] implemented in Seurat, later adapted to *scanpy* [9], or the one implemented in *scry* [10].

A new generation of FS methods emerged when Svensson discovered that the proportion of zeros in droplet-based scRNA-seq data, originally assumed to be dropouts, was tightly related to the mean expression of genes, following a negative binomial (NB) curve [11]. Genes with an expected lower percentage of zeros tend to have an even expression across the entire set of cells. Conversely, genes with a higher than expected percentage of zeros might possess biological relevance because they are expressed in fewer cells than expected, and these cells might be associated to a specific cell type or state.

This finding opened the path for new FS methods that would rely on genes that showed a greater than expected proportion of zeros, according to their mean expression. These methods are based on a null distribution of some property of the dataset, and genes whose behavior differs from the expected are selected. The FS method *nbumi,* a negative binomial method based on *M3Drop* [12], works under this premise. *Nbumi* fits the NB zero-count probability distribution to the dataset, and selects genes of interest calculating *p*-values of observed dropout rates. *M3Drop* works similarly by fitting a Michaelis-Menten model instead of the NB from *nbumi.*

In summary, existing FS methods assume that an unexpected distribution of counts for a particular gene in a dataset is explained by cells belonging to different cell types. However, we observe that there are three main patterns of expression according to the distribution of zeros of a particular gene and overall transcriptional similarity (expression of all genes), as explained in detail in Figure 1: a) a gene evenly expressed across cells, or a gene expressed by a subset of cells, which can be b1) transcriptionally separate or b2) transcriptionally similar. Thus, in some cases a particular gene shows an unexpected distribution of counts because a subset of cells are expressing it but those cells might not be transcriptionally similar.

**Figure 1.**
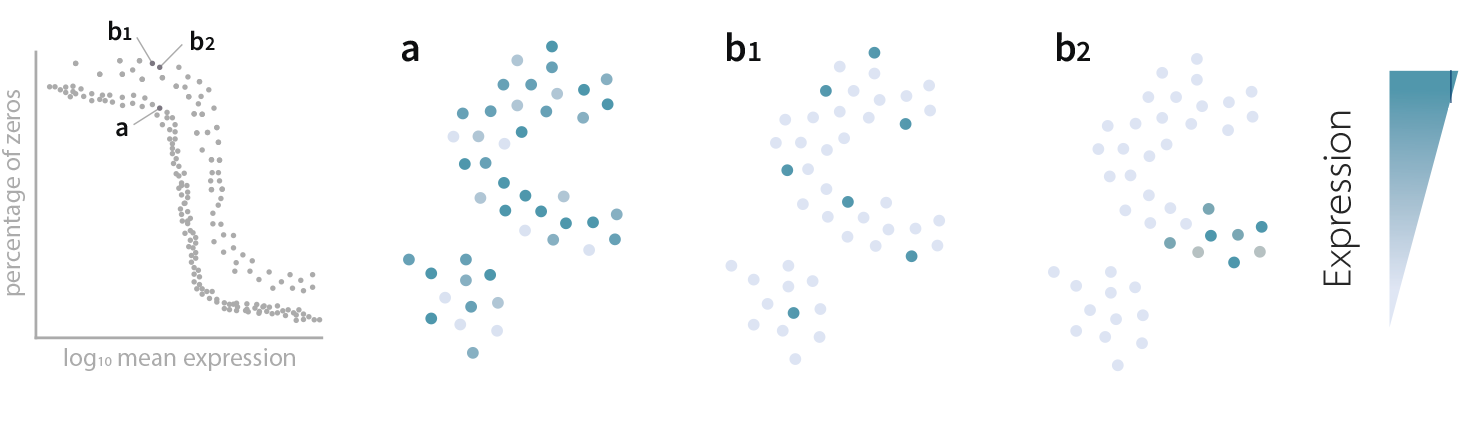
Distribution of gene expression in three scenarios. There are three main patterns of expression for any particular gene in a single-cell dataset: a) The gene is expressed evenly across cells in the dataset, which probably means it does not define any particular cell type. b) A gene shows an unexpected distribution of zeros, because it is only expressed by a subset of cells. Within case b, there are two possible patterns. b1) The gene is highly expressed by a subset of transciptionally different cells (i.e. cells that are not collocalized in the dimensionally reduced map) and b2) the gene is highly expressed by cells that share an overall transcriptomic profile. *Triku* preferentially selects the genes shown in the b2 pattern. When looking at the proportion of zeros, genes in cases b1 and b2 show an increased proportion of zeros with respect to a, but they are indistinguishable from each other by that metric.

Here we present *triku,* a FS method that selects genes that show an unexpected distribution of zero counts and whose expression is localized in cells that are transcriptomically similar. Figure 2 summarizes the feature selection process. *Triku* identifies genes that are locally overexpressed in groups of neighboring cells by inferring the distribution of counts in the vicinity of a cell and computing the expected distribution of counts. Then, the Wasserstein distance between the observed and the expected distributions is computed and genes are ranked according to that distance. Higher distances imply that the gene is locally expressed in a subset of transcriptionally similar cells. Finally, a subset of relevant features is selected using a cutoff value for the distance. *Triku* outperforms other feature selection methods on benchmarking and artificial datasets, using unbiased evaluation metrics such as Normalized Mutual Information (NMI) or Silhouette. Of note, features selected by *triku* are more biologically meaningful.

**Figure 2.**
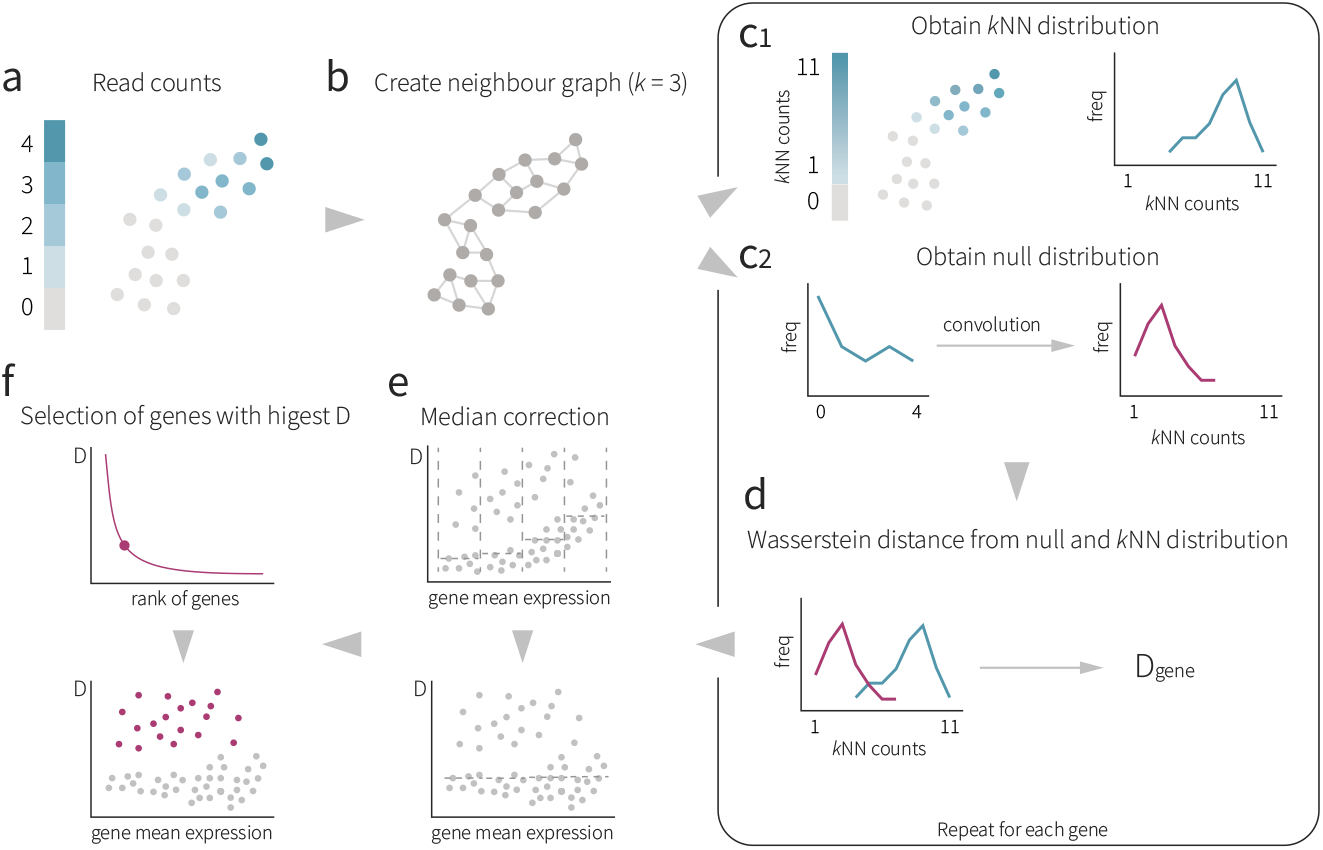
Graphical abstract of *triku* workflow. a) DR representation of the gene expression from the count matrix from a dataset, where each dot represents a cell. b) kNN graph representation with 3 neighbors. For each cell the *k* transcriptomically most similar cells are selected (3 in this example). c1) Considering the graph in b) for each cell with positive expression, the expression of its k neighbors is summed to yield the kNN distribution in blue. c2) With the distribution of reads (blue line), the null distribution is estimated by sampling *k* random cells. d) The null and kNN distributions of each gene are compared using the Wasserstein distance. e) For each gene, its distance is plotted against the log mean expression, and divided into *w* windows (4 in this example). For each window, the median of the distances is calculated and subtracted to the distances in that window. f) All corrected distances are ranked and the cutoff point is selected.

## 2 Results

The objective of FS methods is to select the features that are the most relevant in order to understand and explain the structure of the dataset. In the context of single-cell data, this means finding the subset of genes that, when given as input to a clustering method, will yield a clustering solution where each cluster can be annotated as a putative cell type.

Initially, we generated artificial datasets with the *splatter* package [13], so that cells belonging to the same cluster have a similar gene expression. All datasets contained the same number of genes, cells and populations, but differed in the de.prob parameter value. This parameter was set so that higher values indicate a higher probability of genes being differentially expressed, resulting in more resolved populations. A combination of 8 de.prob values, from 0.0065 to 0.3 were used (see Methods). In addition, we tested *triku* on two biological benchmarking datasets by Ding et al. [14] and Mereu et al. [15] that have been expert-labeled using a semi-supervised procedure. Both benchmarking datasets are composed of individual subsets of data with different library preparation methods (10X, SMART-seq2, etc.) in human Peripheral Blood Mononuclear Cells (PBMCs) (Mereu and Ding) and mouse colon (Mereu) and cortex (Ding) cells.

We have evaluated the relevance of the features selected by *triku* by comparing them to the ones selected using other feature selection methods. The relevance of the features was first measured using metrics associated to the efficacy of clustering, and then using metrics to evaluate the quality of the genes selected.

We made six types of comparisons between the subsets of genes selected by each feature selection method: 1) the ability to recover basic dataset structure (main cell types) in artificial and biological datasets, 2) the ability to obtain transcriptomically distinct cell clusters, 3) the overlap of features between different FS methods, 4) the localized pattern of expression of the features selected, 5) the ability to avoid the overrepresentation of mitochondrial and ribosomal genes and 6) the biological relevance of the genes by studying the composition and quality of the gene ontology (GO) terms obtained.

### 2.1 *triku* efficiently recovers cell populations present in sc-RNAseq datasets

The first set of metrics evaluates the ability to recover the original cell types based on the NMI index, and the cluster separation and cohesion using the Silhouette coefficient.

#### 2.1.1 NMI

NMI measures the correspondence between a labelling considered as the ground truth and the clustering solution that we obtained using the genes selected by *triku* and other FS methods *(scanpy, std, scry, brennecke, m3drop, nbumi).*

First, we evaluated how well the clustering using the genes selected by the FS methods was able to recover the same populations that were defined when generating the artificial datasets. Figure 3 shows that *triku* is among the best three feature selection methods for a wide range of de.prob values. For low values of de.prob –below 0.05–, where the selection of genes that lead to a correct recovery of cell populations is more challenging, *triku* notably outperforms the rest of the FS methods. NMI values obtained with *triku* are 0.1 to 0.2 higher than the second and third best FS methods. In addition, the results obtained when using the first 250 selected genes were comparable to those obtained when selecting 500 genes.

**Figure 3.**
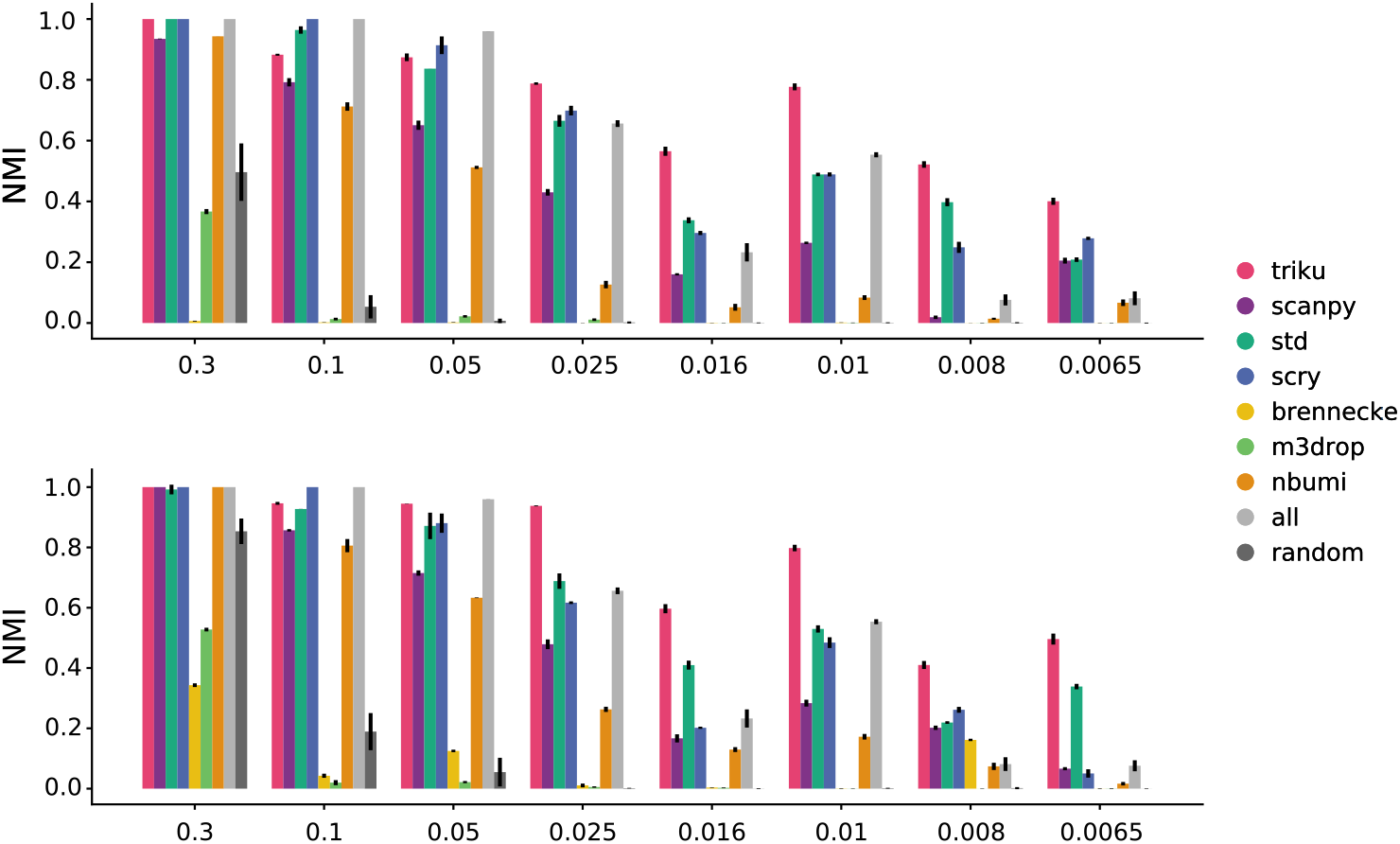
Comparison of NMI for FS methods on artificial datasets. Barplots of the NMI for all FS methods with different artificial datasets, using the top 250 (top) and 500 (bottom) features of each FS method. The probability of the selected genes being differentially expressed between clusters *(de.prob)* is shown in the X axis. Higher NMI values mean better recovery of the cell populations. Note that in category *all,* all features are selected, not the top 250 or 500, therefore their NMI values are the same in both graphs.

We also studied how well the genes selected led to a clustering solution that was similar to the manually-assigned cell labels in the biological benchmarking datasets, as shown in Figure 4. For each dataset, the variability between NMI scores was quite low, meaning that features selected with the different methods yielded clustering solutions that were quite similar to the manually-labeled cell types, although there are some exceptions to this rule–e.g. Brennecke in Ding datasets, which showed notably reduced NMI values–. In some datasets, for instance, 10X human, QUARTZseq human and SMARTseq2 human from Mereu’s benchmarking set, features selected by FS methods did not lead to increased NMI values as compared with randomly selected genes.

**Figure 4.**
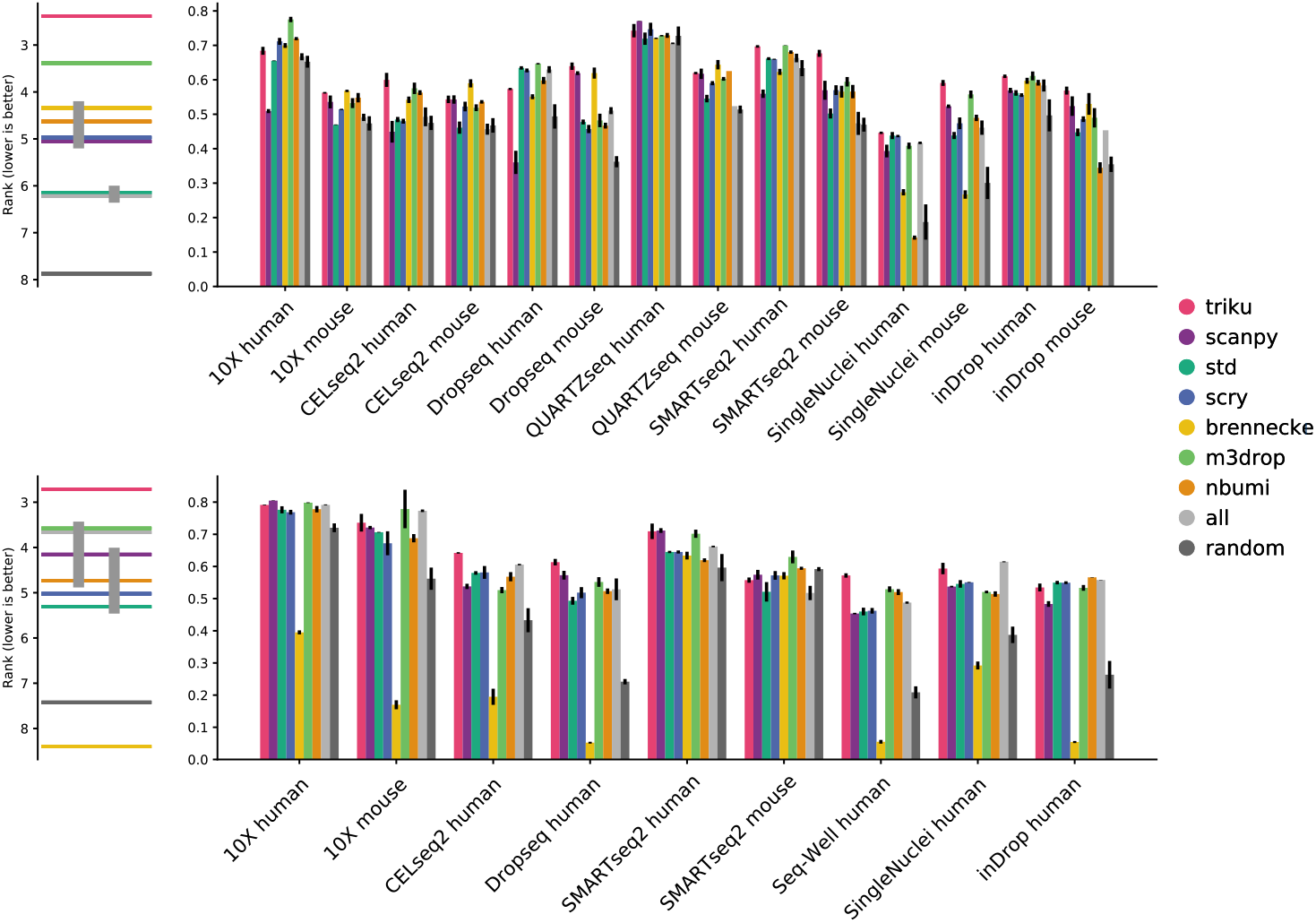
NMI for annotated cell types in Mereu and Ding datasets. Barplots of Silhouette coefficient for Mereu (top) and Ding (bottom) datasets. Each barplot represents the mean over 5 runs, and the vertical bar is the standard deviation. The plot on the left is a critical difference diagram, where each horizontal bar represents the mean rank for all datasets. If two or more bars are linked by a vertical bar, the mean ranks for those FS methods are not significantly different (Quade test, *α =* 0.05).

Despite the differences in NMI between methods being small for each particular dataset, post-hoc analysis revealed that *triku* is significantly the best ranked method across all datasets. To do the post-hoc analysis, we ranked for each dataset the NMI of each FS method. Figure 4 (left) shows the mean rank of each FS method across datasets. *Triku* is the best-ranked FS method in both Mereu’s and Ding’s benchmarking datasets, with a mean rank of 2.7 and 2.8, respectively. *M3drop* is the second best-ranked FS method and *triku* is in both cases statistically significantly better (Quade test, *p* < 0.05).

#### 2.1.2 Silhouette coefficient

Another important aspect of the genes selected by FS methods in scRNA-seq data analysis is their ability to cluster data into well-separated groups that are transcriptomically similar. We used the Silhouette coefficient to measure the compactness and separation-degree of cell communities obtained with a clustering method. When the same clustering algorithm is used on a dataset but using two different FS methods, the differences in the resulting Silhouette coefficients can be entirely attributed to the features selected by those methods. We assume that FS methods that increase the separation between clusters and the compactness within clusters are better at recovering the cell types present in the dataset.

Figure 5 shows the Silhouette coefficients obtained with the different FS methods. For the Mereu and Ding datasets, we observed that *triku* was the best-ranked method–mean rank of 1.8 and 1.1–, and the second best-ranked methods were *m3drop* and *scanpy* with a mean rank of 3.8 and 2.2, respectively. In both cases, the difference between *triku* and the second-ranked method was statistically significant (Quade test, *p* < 0.05).

**Figure 5.**
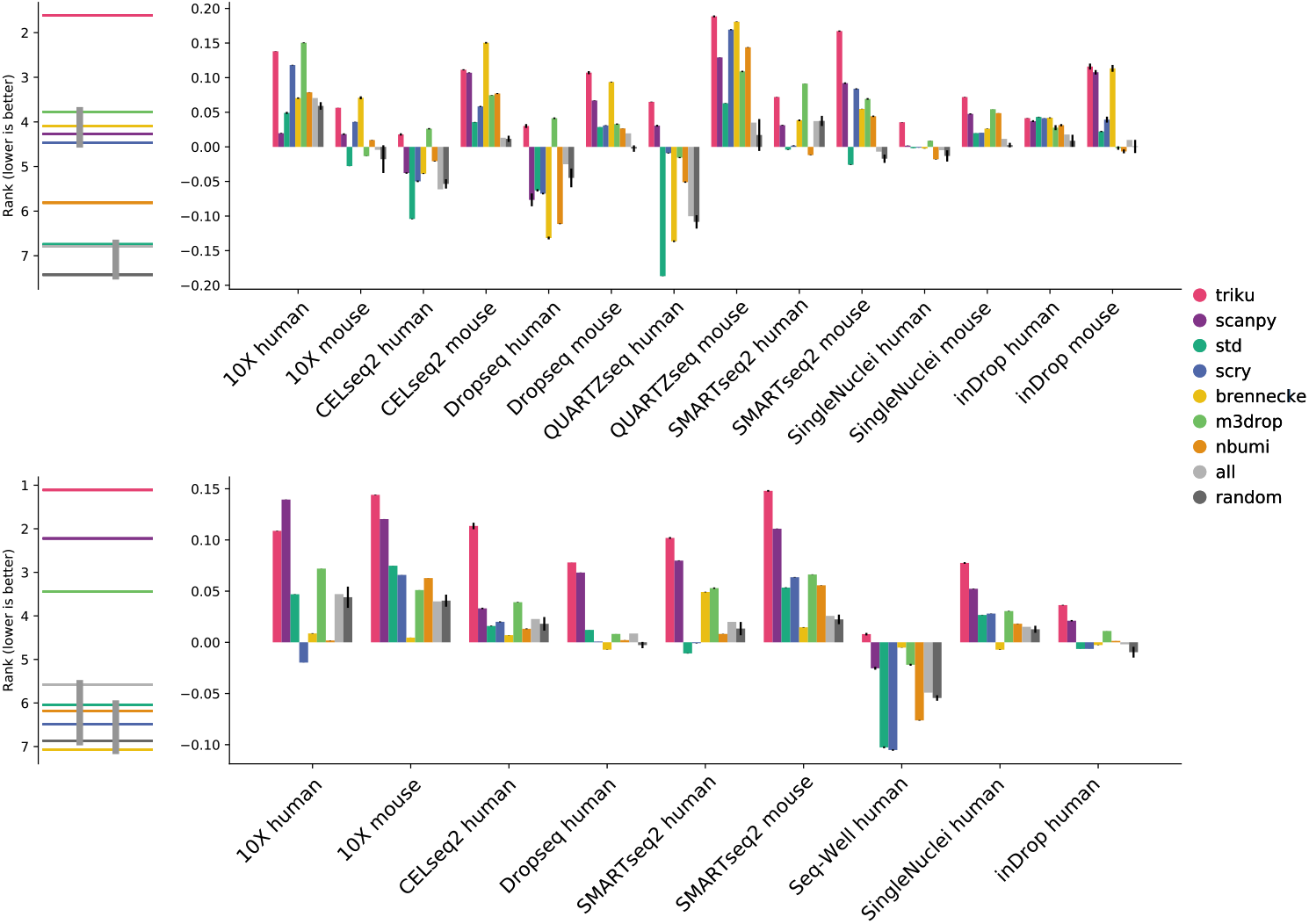
Silhouette coefficients for annotated cell types in Mereu and Ding datasets. Barplots of Silhouette coefficient for Mereu (top) and Ding (bottom) datasets. Each barplot represents the mean of 5 seeds, and the vertical bar is the standard deviation. The plot on the left is a critical difference diagram, where each horizontal bar represents the mean rank for all datasets and all seeds. If two or more bars are linked by a vertical bar, the mean ranks for those FS methods are not significantly different (Quade test, *α* = 0.05).

We performed an additional analysis using the labels obtained with leiden clustering instead of the manually curated cell types (Figure S1). Again, *triku* outperformed the rest of the FS methods showing a statistically significant best mean-rank.

### 2.2 Genes selected by different FS methods show limited overlap

Next, we studied the characteristics of the genes selected by *triku* and compared them to the genes selected by other methods.

Initially, we studied the level of consistency between the results obtained using different FS methods by studying their degree of overlap, as shown in Figure 6. In order to compare between equally sized gene lists, we ranked the genes based on p-values or scoring value from each FS method and set the number of genes selected by *triku* as a cutoff to select the first genes. Although the genes selected by the different methods yielded clustering solutions that are highly consistent, as shown in the previous section, we did not see any clear gene overlap pattern between pairs of FS methods. In fact, there is no correlation between the degree of overlap between the genes selected by the different methods and the clustering solutions that are obtained when using those genes as input.

**Figure 6.**
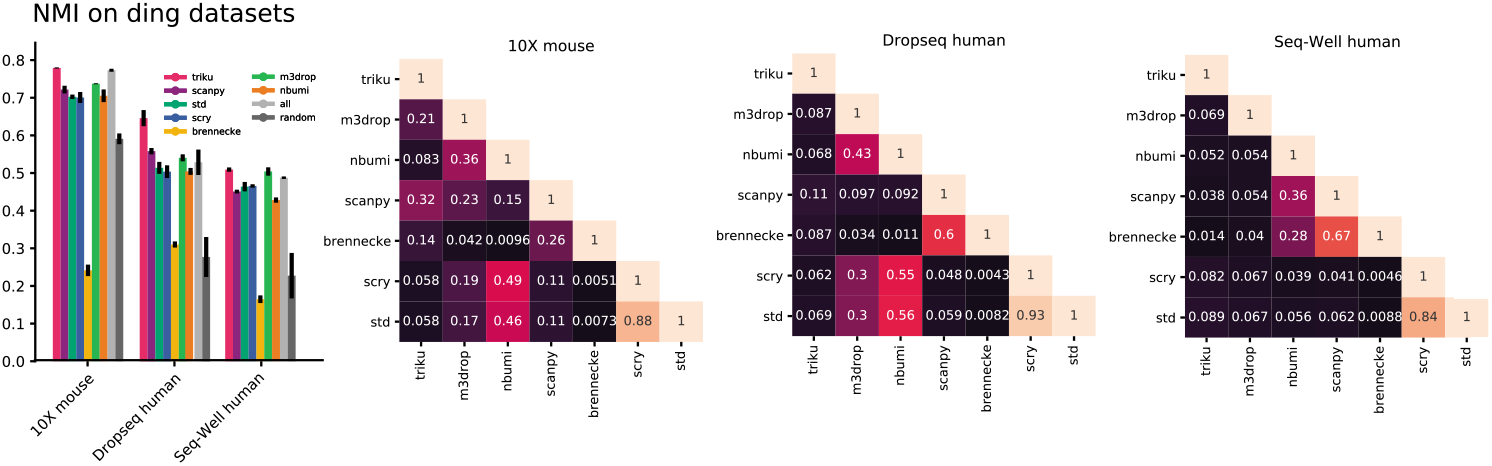
Heatmaps of overlap of features between pairs of methods. For each pair of methods, the value represents the proportion of features that are shared between the two methods. The number of genes selected in each method is the automatic cutoff by *triku.*

For instance, we found an overlap of 11% between the genes selected by *scanpy* and *std* for the 10x mouse dataset, yet the NMI between the clustering solutions obtained with each of them and the expert-labeled cell types was 0.7. On the other hand, the overlap between *scanpy* and *Brennecke* is one of the highest across datasets (ranging from 26 to 67%), yet the differences between their corresponding NMI scores are 0.45.

### 2.3 *triku* selects genes that are biologically relevant

Based on these results, we studied the biological relevance of the genes selected by different FS methods in three alternative ways.

Genes whose expression, or lack thereof, is limited to a single population are more likely to be cell-type specific and thus might be better candidates as positive or negative cell population markers. Therefore, we studied which are the best FS methods to select genes showing a localized expression pattern.

Mitochondrial and ribosomal genes are usually highly expressed and many FS methods tend to overselect them despite them not being particularly relevant in most single-cell studies and are commonly excluded from downstream analysis [16, 17, 18]. Assuming that in these benchmarking datasets ribosomal and mitochondrial genes are not as relevant to the biology of the dataset, we measured the percentage of these genes in the subset of genes selected by *triku* and compared it to other FS methods.

Lastly, we analyzed the biological pertinence of the selected genes by performing Gene Ontology Enrichment Analysis (GOEA) on a dataset of immune cell populations whose underlying biology is well understood, as a robust indicator of FS quality.

#### 2.3.1 Selection of locally-expressed genes

We first studied the expression pattern of genes selected by *triku* and other methods, as shown in Figure S2. We observed that out of the 9 populations of the artificial dataset, when a gene is selected by *triku*–exclusively or together with other FS methods–, one of the populations had a markedly higher or lower expression compared to the rest. On the other hand, when a gene is selected by other FS methods and not by *triku*, we do not observe any population-specific expression pattern. For instance, genes exclusively selected by *scanpy* had a wide expression variation across clusters, but they were not exclusive of one or two clusters. Features selected by *std* and *scry* showed some variation, but it was overshadowed by the high expression of the gene, and therefore not relevant under the previous premise.

To evaluate the cluster expression of selected genes in benchmarking datasets, for each gene we scaled its expression to the 0-1 range, and sorted the clusters so that the first one had the greatest expression. Figure S3 shows the expression patterns for several benchmarking datasets. We see that, in most datasets, *triku* showed more biased expression patterns, that is, genes selected by *triku* were expressed, on average, on fewer clusters than the genes selected by other FS methods. The second and third best methods were *scanpy* and *brennecke,* with similar or slightly less biased expression patterns as compared to *triku*. With these methods, up to 80% of the expression of the gene was usually restricted to the 2 to 3 clusters that most express it.

*m3drop* and *nbumi* performed similarly, and showed an expression distribution across clusters similar to a random selection of genes, which was slightly biased towards 3 to 5 clusters accumulating up to 80 % of the expression of the gene. Lastly, *std* and *scry* methods were the least biased, and showed almost a linear decrease of expression percentage across clusters, with 4 to 6 clusters accumulating up to 80 % of the expression of the gene.

#### 2.3.2 Avoidance of mitochondrial and ribosomal genes

Table 1 shows the percentage of genes that code for ribosomal and mitochondrial proteins within the genes selected by different FS methods in the two sets of benchmarking datasets. We observed that *std* and *scry* were the only methods that tended to overselect mitochondrial and ribosomal genes. Among the rest of the methods, *triku* showed percentages that were comparable to the rest of the methods, and slightly lower for the Ding datasets.

**Table 1.**
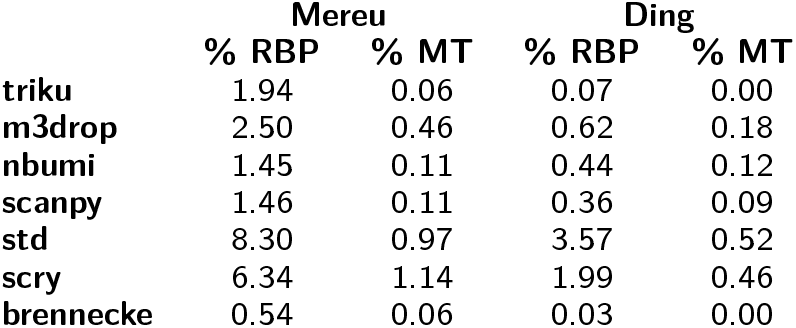
Percentage of ribosomal protein (RBP) and mitochondrial (MT) genes appearing within the selected genes by each FS method.

#### 2.3.3 Selection of genes based on gene ontologies

We assessed the quality of the GO output by studying its term composition. We selected two PBMC datasets from the Ding datasets: the 10X human and the Dropseq human. We used PBMC datasets for this analysis because their cell-to-cell variability has been extensively studied using single-cell technologies as Fluorescence Activated Cell Sorting (FACS) and scRNA-seq [19, 20, 21, 22, 23]. Using these datasets, we measured the proportion of GO terms obtained in the output that were tightly related to the biological system under study.

Figures 7 and S4 show the first 25 GO terms obtained with the genes selected by each FS method on the two PBMC datasets, where the terms tightly related to immune processes–chosen by three independent assessors–have been highlighted. We observed that *triku* was the FS method that yielded the most terms directly related to immune processes, with 23/25 and 19/25 related terms in the Ding Dropseq and 10X datasets, respectively. Examples of terms that we considered to be tightly related to immune processes included *B cell receptor signalling pathway, neutrophil degranulation* and *T cell proliferation.* The next methods were *scanpy* and *m3drop,* whose performances were comparable to that of *triku* for the 10X dataset (23/25) but less robust for the Dropseq dataset (10/25 and 9/25 related terms). The rest of the FS methods mainly selected genes that were related to general cell functions such as RNA processing, protein processing and cell-cycle regulation.

**Figure 7.**
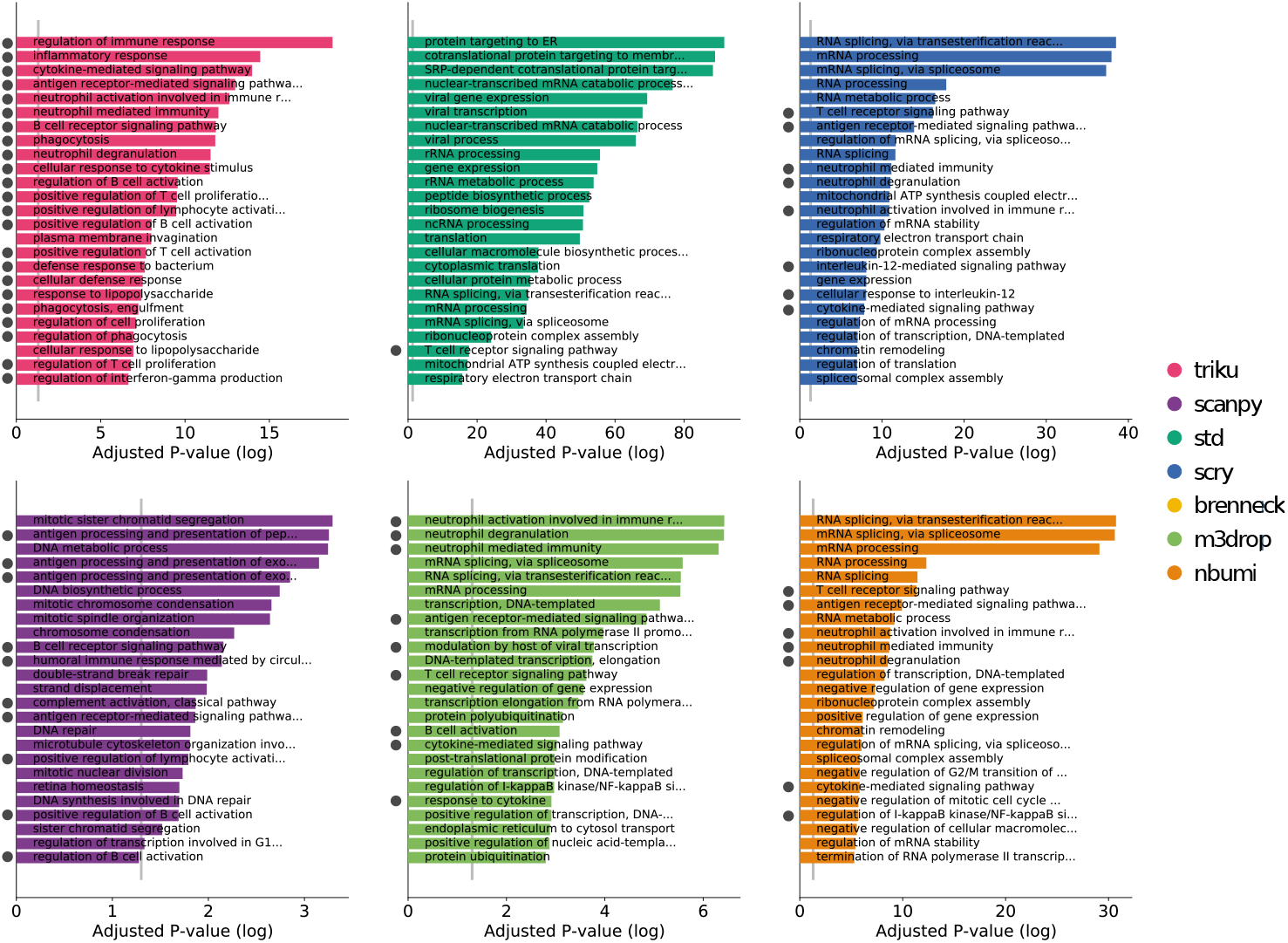
Barplot of p-values of GOEA. Each bin represents the number of features selected for each method, in Mereu et al. mouse Dropseq dataset. The y value is the −log10 adjusted p-value for the best 25 ontologies. On the bottom, the bar plot shows the names of the ontology terms for the case with the best 1000 features. In immune datasets, gray dots at the left of each term represent that that term is directly-related to an immune process. Non-dotted terms refer to more general processes that may or may not be related to immune processes.

## 3 Discussion

FS methods are a key step in any scRNA-seq sequencing analysis pipeline as they help us obtain a dimensionally reduced version of the dataset that captures the most relevant information and eases the interpretation and understanding of its underlying biology. However, every FS method relies on a set of assumptions regarding what characteristics make a gene relevant. FS methods that sort genes according to their dispersion assume that gene expression variability is indicative of its biological relevance. FS methods like *nbumi* and *m3drop* assume that genes showing a proportion of zero-counts that is greater than expected (according to a null distribution) are more likely to be informative. *Triku* assumes that genes that have a localized expression in a subset of cells that share an overall transcriptomic similarity are more likely to define cell types. A general trend in FS method design has been to refine the requirements that a gene must meet in order for it to be selected, from the more general dispersion-based to more sophisticated formulations. It is noteworthy that the requirements in *triku* are consistent with the previous dispersion-based and zero-count-based formulations, but involve a new aspect that we consider essential for an accurate gene selection: a localized expression in neighboring cells. Another important advantage of *triku* over FS methods that consider the zero-count distribution is that, unlike *m3drop* and *nbumi*, *triku* does not assume gene counts to follow any particular distribution, since it estimates the null distribution from the dataset, thus extending the range of single-cell technologies that it can use beyond droplet-based technologies.

We verified the locality of the genes selected by *triku* in different artificial and real scRNA-seq datasests and concluded that, on average, the expression of *triku*-selected genes is restricted to fewer, well-defined clusters. In addition, the clusters obtained when using *triku*-selected genes as input for unsupervised clustering in both artificially generated and biological datasets have a better resolved pattern structure, as shown by their increased Silhouette coefficients. In the case of artificial datasets, where the degree of mixture between clusters can be predefined, *triku* proved to be able to recover the originally-defined cell populations. In fact, we found that the higher the degree of mixture between clusters, the more obvious the advantage of *triku* over the rest of the FS methods tested.

An important difficulty in the interpretation of single-cell data is that we must consider that cell-to-cell variability has both technical and biological components. I.e., it is difficult to know whether a set of genes is differentially expressed between cell clusters due to technical reasons (differences in the efficiency of mRNA capture, amplification and sequencing) or if it constitutes a biological signal. Moreover, there is a wide range of sources of biological variability within a dataset, some of which might not be of interest depending on the experimental context. For instance, fluctuations in genes that regulate the cell cycle constitute a source of biological variability that is often disregarded. This has been extensively studied and addressed in a number of ways: normalization, regression of unwanted sources of variation, etc. [24, 25, 26, 27].

The expression of genes whose variability is associated with technical reasons tend to have a high dispersion but their expression is usually not restricted to a few clusters. A good example of these genes are the ribosomal and mitochondrial genes, which are expressed across all cell types at different levels. Our results show that these genes are in fact selected by the majority of compared FS methods due to their high expression and cell-to-cell variability, but are less likely to be selected by *triku*, since they do not usually meet the locality requirement. Additionally, when performing GOEA, we observed that the list of genes obtained with *triku* were more enriched for terms that are specifically related to a biological process of the system under study.

In our work, we have observed that the genes selected by different FS methods might show little overlap between them. This phenomenon has been described elsewhere [28]. In fact, gene covariation and redundancy is a well characterized effect that has been observed in omics studies. The effect of redundancy arises from the fact that different cell types must have a common large set of pathways to be active. The difference between cell type and cell state is that two cell types might have large sets of pathways that are different between each other, and two cell states will only differ in a few pathways. Since pathways are composed of many genes, only choosing a reduced set of genes from a set of pathways from cell type A and B might be enough to differentiate them, and we might not need to select all genes from all pathways. This “paradigm” explains several effects. Qiu et al. described that scRNA-seq datasets could preserve basic structure after gene expression binarization [29] or by conducting very shallow sequencing experiments [4]. This can be explained by the fact that only a few genes are necessary to describe the main cell populations in a single-cell dataset, and the presence/absence of a certain marker is often more informative than its expression level. This is related to the notion that despite the high dimensionality of omics studies, most biological systems can be explained in a reduced number of dimensions. Moreover, some authors have claimed this low dimensionality to be a natural and fundamental property of gene expression data [4]. This highlights the importance of designing accurate FS methods that extract the fundamental information from single-cell datasets.

*Triku* Python package is available at https://gitlab.com/alexmascension/triku and can be downloaded using PyPI. *Triku* has been designed to be compatible with *scanpy* syntax, so that *scanpy* users can easily include *triku* into their pipelines.

## 4 Methods

The *triku* workflow is further described in Suplementary Methods.

### 4.1 Artificial and benchmarking datasets

In order to perform the evaluation of the FS methods we used a set of artificial and biological benchmarking datasets.

Artificial datasets were constructed using *splatter* R package (v 1.10.1). Each dataset contains 10,000 cells and 15,000 genes, and consists of 9 populations with abundances in the dataset of {25%, 20%, 15%, 10%, 10%, 7%, 5.5%, 4%, 3.5%} of the cells. Each dataset contains a parameter, de.prob, that controls the probability that a gene is differentially expressed. Lower de.prob values (< 0.05) imply that different populations have fewer differentially expressed genes between them and, therefore, are more difficult to be differentiated. Selected values of de.prob are {0.0065, 0.008, 0.01, 0.016, 0.025, 0.05, 0.1, 0.3}. Populations in datasets with de.prob values above 0.05 are completely separated in the low-dimensionality representation with UMAP, even without feature selection (Figure S5).

Regarding biological datasets, two benchmarking datasets have been recently published by Mereu et al. [15] and Ding et al. [14]. The aim of these two works is to analyze the diversity of library preparation methods, e.g. 10X, SMART-seq2, CEL-seq2, single nucleus or inDrop. Mereu et al. use mouse colon cells and human PBMCs to perform the benchmarking, whereas Ding et al. use mouse cortex and human PBMCs. There are a total of 14 datasets in Mereu et al. and 9 in Ding et al. An additional characteristic of these datasets is that they have been manually annotated, and this annotation is useful as a *semi* ground truth. Ding dataset files were downloaded from Single Cell Portal (accession numbers SCP424 and SCP425), and cell type metadata is located within the downloaded files. Mereu datasets were downloaded from GEO database (accession GSE133549), and cell type metadata was obtained under personal request.

### 4.2 FS methods

*Triku* is compared to the following FS methods:

- Standard deviation (*std*). Computed directly using Numpy (v 1.18.3).
- *brennecke* [7]: fits a curve based on the square of the coefficient of variation *(CV*^2^) versus the mean expression (μ) of each gene and selects the features with higher CV^2^ and μ. The features are selected with the *BrenneckeGetVariableGenes* function from M3Drop R package (v 1.12.0).
- *scry* [10]: computes a deviance statistic for counts based on a multinomial model that assumes each feature has a constant rate. The features are selected with the *devianceFeatureSelection* function from scry R package (v 0.99.0).
- *scanpy* [9]: selects features based on a z-scored deviation, adapted from *Seurats* method. The features are selected with the *sc.pp.highly_variable_genes* function from *scanpy* (v 1.6.0).
- *M3Drop* [12]: fits a Michaelis-Menten equation to the percentage of zeros versus μ, and selects features with higher percentages of zeros than expected. The features are selected with the *M3DropFeatureSelection* function from M3Drop R package.
- *nbumi*: it acts in the same manner as *M3Drop*, but fitting a negative binomial equation instead of a Michaelis-Menten equation. The features are selected with the *NBumiFeatureSelectionCombinedDrop* function.

#### 4.2.1 FS and dataset preprocessing

To make the comparison between FS methods, each feature is ranked based on the score provided by each FS method. Calculating the ranking instead of just selecting the features allow us to select different numbers of features when needed. By default, the number of features is the one automatically selected by *triku*. Additionally, in some contexts, analyses are performed with all features or with a random selection of features.

After the ranking of genes is computed, dataset processing is performed equally for all methods, in artificial and benchmarking datasets. Datasets are first log transformed –if required by the method–, and PCA with 30 components is calculated. Then, the *k*-Nearest Neighbors (*k*NN) matrix is computed setting *k* as 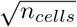. Uniform Manifold Approximation and Projection (UMAP) (v 0.3.10) is then applied to reduce the dimensionality for plotting. If community detection is required, leiden (v 0.7.0) is applied selecting the resolution that matches the number of cell types manually annotated in the dataset. This procedure is repeated with 10 different seeds. This conditions the output of *triku*, random FS, PCA projection, neighbor graph, leiden community detection, and UMAP.

#### 4.2.2 NMI calculation in artificial and benchmarking datasets

In order to compare the leiden community detection results with the ground-truth labels from artificial and biological datasets, we used the Normalized Mutual Information (NMI) score [30].

If *T* and *L* are the labels of the cell types (true populations) and leiden communities respectively, the NMI between *T* and *L* is:

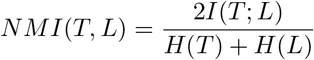

Where *H*(*X*) is the entropy of the labels, and *I*(*T*; *L*) is the mutual information between the two sets of labels. This value is further described in [31]. We used scikit-learn (v 0.23.1) implementation of NMI, *sklearn.metrics.adjusted_mutual_info_score.*

One of the advantages of NMI against other mutual information methods is that it performs better with label sets with class imbalance, which are common in singlecell datasets, where there are differences in the abundance of cell types.

On artificial datasets, leiden was applied using the first 250 and 500 selected features, and the resulting community labels were compared with the population labels from the dataset. On benchmarking datasets, leiden was applied with the manually-curated cell types.

#### 4.2.3 Silhouette coefficient in benchmarking datasets

In order to assess the clustering performance of the communities obtained with benchmarking datasets we used the Silhouette coefficient. The Silhouette coefficient compares the distances of the cells within each cluster (intra-cluster) and between clusters (inter-cluster) within a measurable space. The distance between two cells is the cosine distance between their gene expression vectors, considering only the genes selected by each FS method. The greater the distance between cells that belong to different clusters and the smaller the distance between cells from different cluster, the greater the Silhouette score.

In order to calculate the Silhouette coefficient for a cell *c* within cluster *C_i_* (out of *n* clusters), the mean distance between the cell and the rest of the cells within the cluster is computed using the gene expression:

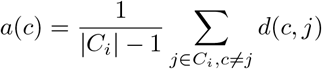

Then, the minimum mean distance between that cell and the rest of cells from other clusters is computed:

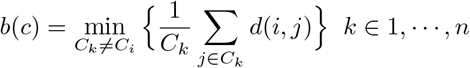

Then the Silhouette coefficient is computed as

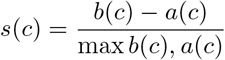

Higher Silhouette scores imply a better separation between clusters and, therefore, a better performance of the FS method. We used scikit-learn implementation of Silhouette, *sklearn.metrics.silhouette_score.*

#### 4.2.4 Overlap between gene lists

In order to calculate the overlap between selected features for each FS method, we applied the Jaccard index [32]: 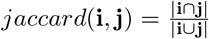, where **i**, **j** are the sets of genes selected by the two FS methods.

#### 4.2.5 Performance of gene selection and locality measures

In order to assess the performance of different FS methods selecting genes that are relevant for the dataset, we applied two different strategies for artificial and biological datasets.

For artificial datasets, we selected 4 representative genes of each of the combinations of genes shown in Figure S2. Then we calculated the mean expression of each of the for genes in each population, and we represent this information in the barplots.

For benchmarking datasets, in order to represent the Figure S3, for each dataset and FS method we used the following procedure: for each gene, the expression was scaled to sum 1 across all cells. Then, leiden clustering was run with resolution parameter value 1.2. For each cluster, the proportion of the expression was calculated, and the clusters were ordered so that the first cluster is the one that concentrates the majority of the expression. To create Figure S3, the average value of the proportion of expression is calculated.

#### 4.2.6 Proportion of ribosomal and mitochondrial genes

When calculating the proportion of mitochondrial and ribosomal genes, the list of existing ribosomal and mitochondrial proteins was calculated by extracting the genes starting with *RPS, RPL* or *MT-.* The proportion of mitochondrial or ribosomal genes is the quotient between the genes of the previous list that appear selected by that FS method, and the genes in the list.

#### 4.2.7 GO enrichment analysis

In order to calculate the sets of gene ontologies enriched for the selected features of each FS method, we used python *gseapy* (v 0.9.17) module *gseapy.enrichr* function with the list of the first 1000 selected features against the *GO_Biological_Process_2018* ontology. From the list of enriched ontologies, the 25 with the smallest adjusted *p*-value were selected.

#### 4.2.8 Ranking and CD

During calculation of NMI and Silhouette coefficients, to evaluate the overall performance of the FS methods across different datasets, the FS methods are ranked –where 1 is the best rank–. The methodology proposed by Demšar [33] is used to test for significant differences among FS methods in the datasets: The Friedman rank test is applied to test whether the mean rank values for all FS methods are similar (null hypothesis). If the Friedman rank test rejects the null hypothesis (α < 0.05), this implies a statistically significant difference among at least two FS methods. If the null hypothesis is refuted we apply the Quade post-hoc test between all pairs of FS methods to check which pairs of FS methods are significantly different (α < 0.05). These results are then plotted in a critical difference diagram.

## 5 Abbreviations

scRNA-seq: Single-cell RNA sequencing
FS: Feature Selection
FE: Feature Extraction
PCA: Principal Component Analysis
NB: Negative Binomial
(NMI): Normalized Mutual Information
FACS: Fluorescence Activated Cell Sorting
GO: Gene Ontology
GOEA: Gene Ontology Enrichment Analysis
PBMC: Peripheral Blood Mononuclear Cells
UMAP: Uniform Manifold Approximation and Projection
kNN: *k*-Nearest Neighbors

## Declarations

### Ethics approval and consent to participate

Not applicable.

### Consent for publication

Not applicable.

### Availability of data and software

Ding dataset files were downloaded from Single Cell Portal (accession numbers SCP424 and SCP425), and cell type metadata is located within the downloaded files. Mereu datasets were downloaded from GEO database (accession GSE133549), and cell type metadata was obtained under personal request. Triku software and analysis notebooks are available at https://www.gitlab.com/alexmascension/triku.

### Author’s contributions

Conceptualization: AMA; Funding Acquisition: MJA-B, AMA, OI-S; Investigation: AMA, OI-S, MJA-B, AI; Methodology: AMA, OI-S, II; Project Administration: AI, MJA-B; Resources: MJA-B; Software: AMA, OI-S; Supervision: II, AI, MJA-B; Visualization: AMA, OI-S; Writing - Original Draft Preparation: AMA, OI-S; Writing - Review and Editing: AMA, OI-S, II, MJA-B, AI.

## Acknowledgements

We would like to thank Amaia Elícegui, Ainhoa Irastorza and Paula Vázquez for the assessment of the immune Gene Ontology terms.

