## Supplementary Materials for "Triku: a feature selection method based on nearest neighbors for single-cell data"

February 12, 2021

#### 1 Supplementary Results

##### 1.1 *Triku* workflow

The premise of *triku* is that, for genes with similar expression levels, the expression pattern can be categorized in three states (Figure 1). On the one hand, if the gene is expressed throughout the cells with similar expression levels it does not provide any useful information about specific cell types associated to that gene (a). On the other hand, the expression of the gene can be localized in a subset of cells. This subset can be of transcriptomically different cells (b1), or transcriptomically similar cells (b2). The latter case is interesting because these cells, sharing similar transcriptomes, will have high expression of other genes, and therefore a putative biological function. *Triku* aims to select genes of case (b2) while avoiding the selection of genes of case (a) and (b1).

In the Figure 2 we show the workflow followed by *triku* to select features of the type (b2). It accepts as input raw or log-transformed read count matrices, and assigns a score of goodness to each gene. To compute that score, the first step is to create a neighbour graph. The graph is learned by selecting the $k$ cells with the most similar transcriptome to each cell in the dataset–$k$ Nearest Neighbours, or $k$NN for short–. $k$ is arbitrary, and can be fixed by the user, with typical values $\sim \sqrt{n_{cells}}$. Before computing the graph the dimensionality of the dataset is reduced using Principal Component Analysis (PCA) to reduce the computation time of the graph.

Once the $k$NN graph is defined, the distribution of the $k$NN counts for each gene is obtained by summing the counts of each cell and its neighbors for each cell with positive expression in the dataset. Cells that have zero expression are not considered, even if their neighbours have positive expression. The reason for this will be discussed later.

Once the $k$NN distribution is learned for each gene, the next step is to simulate, also for each gene, a null distribution. The null distribution is built by considering $k$ random cells instead of $k$NN ones. This is mathematically translated as applying $k$ times the convolution of the distribution of reads–including zeros–to the distribution of reads without considering zeros.

By applying the convolution of one distribution to another the two probability distributions are mathematically "merged". If the convolution between the distribution of reads without zeros (to simulate the initial choice of cells that express the gene) and the distribution of reads with all cells is applied, the resulting distribution is the one by choosing two cells at random–one with positive expression–and summing up their reads. After applying the convolution $k$ times, the null distribution by choosing one cell with positive expression and $k$ cells at random is created.

The next step of the process is to compare, for each gene, the $k$NN distribution with its corresponding null distribution. Biologically relevant genes have heavily right-tailed $k$NN distributions, and their null distributions are expected to be symmetric; whereas $k$NN distributions of noisy genes will be similar to their associated null distributions (Figure S6).

To quantify the difference between the $k$NN distribution and the null distribution of a gene, the Wasserstein distance between both distributions is computed. The range of Wasserstein distance is in $[0, \infty)$, and the higher the distance the more separated the $k$NN distribution from its null distribution will be.

Once Wasserstein distances have been calculated for all genes, the next step is to apply a normalization procedure based on the mean expression. In some cases, genes with higher expression tend to have higher basal Wasserstein distances, and this bias should be corrected. To correct this, we set a median correction based on a number of windows, $w$. The range of expression–in log scale–is divided in $w$ windows, and for each window, the median distance value of the genes in that window is subtracted to the Wasserstein distance of each gene. After this correction, most genes have a Wasserstein distance near zero, and few have higher distance values, which are the ones to be selected.

The last step is to select the features with highest Wasserstein distance. If the number of features is not defined, a simplification of the *needle method* [1] is applied (described in the methods section) to select the cutoff distance value automatically.

##### Why cells with zero counts are not considered to build the $k$NN count distribution

When calculating the $k$NN and null distributions, all cells could be taken into account instead of just the ones with positive expression. However, with this approach, many of the $k$NN counts will be simply zero, or near zero, for the $k$NN distribution and the null distribution. Although still functional, this drastically decreases the Wasserstein distance of the gene because both distributions are mostly composed of zeros.

An example can be seen in Figure S7, where genes that are appropriate candidates of being selected as relevant show their distance drastically decreased, whereas genes that are not relevant, either because they have higher mean expression values or are noisy, show fewer zero-expressing $k$NN cells, and their Wasserstein distances are similar.

#### 1.2 Robustness of *triku* parameters

There are some parameters in *triku* that affect how features are selected. These parameters are the number of components in PCA ($p$)–to create the $k$NN matrix–, the number of neighbors ($k$), and the number of windows for median correction ($w$). Additionally, some parts in the pipeline are randomized, and random effects have to be considered with different seeds. Each of these parameters can be set by the user, and they also have default values–$p = 30, k = \sqrt{n_{cells}}, w = 100$–.

To check their robustness, *triku* is run for each parameter with a set of values, and the overlap of the first 500, 1500 and 2500 features between the default parameter value and the rest of values in the set is computed. For example, to check for PCA robustness, the overlap of selected features between $p = 30$ (default) and $p \in \{3, 5, 10, 20, 30, 40, 50, 100\}$ is tested. The overlap between the top X features for two $p$ values is defined as the quotient between the number of shared features by X. For each pair of values, the overlap between features is computed varying the same set of seed values, to account for random effects of FS.

Figure S8 shows the analysis of robustness for the 3 parameters, in Ding et al.[2] and Mereu et al.[3] benchmarking datasets. Results are similar for both datasets. The overall trend is that, with the exception of CELseq2, the rest of library preparation methods show good robustness values for $p$, $k$ and $w$. $p$ shows overlap higher than 0.8 for values between 20 and 40, and higher than 0.7 for values between 10 and 50, which are the expected number of components for PCA. Overlap decreases for extreme values like 3 or 5, but those values are not expected to be selected, as they are not standard in single-cell analysis.

Similar results occur for $k$, which is set in $\{\sqrt{N}/20, \sqrt{N}/10, \sqrt{N}/5, \sqrt{N}/2,$ $\sqrt{N}, 1.5\sqrt{N}, 2\sqrt{N}, 4\sqrt{N}\}$. Values between $\sqrt{N}/2$ and $2\sqrt{N}$ show overlap higher than 0.8; $\sqrt{N}/5$ shows overlap values higher than 0.65 for most datasets, which is reasonable considering the vast difference in $k$ to create the $k$NN graph. Extremely low values like $\sqrt{N}/10$ or $\sqrt{N}/20$ show even lower feature overlaps. This behaviour is expected because these $k$ values mean $k = 2$ if the dataset contains 1000 cells, and $k = 10$ if it contains 50000 cells, which are extremely low in comparison with the number of cells in the dataset. $k$NN graphs with these extremely low $k$ values barely capture any global structure, which is required to some extent to find relationships between different cell types. Lastly, the selection of the number of windows is notably robust, with values higher than 0.9 for most datasets, across all values in the number of windows. The tested values of $w$ are $\{10, 20, 30, 50, 100, 200, 500, 1000\}$

Lastly, the effect of seed is analyzed for default values of $p$, $k$, and $w$. This effect is directly observed when the overlap with the default value and itself is computed. Overlap values in most datasets are higher than 0.9, therefore seed is not a key factor when selecting features with *triku*. Therefore, default $p = 30$, $k = \sqrt{n_{cells}}$ values,commonly used in other single-cell pipelines, are robust to changes within reasonable value intervals. The recommended values are $p \in [20 - 50]$ and $k \in [\sqrt{N}/5 - 2\sqrt{N}]$. $w$ is robust overall, and values of $w \in [30 - 500]$ yield almost-identical results.

### 2 Supplementary Methods

#### 2.1 Triku workflow

##### 2.1.1 Calculation of $k$NN counts and null distributions per gene

*Triku* selects genes based on the similarity of cells. A gene is a good candidate if its expression is localized in a set of cells that are transcriptomically similar to each other. On the other hand, a gene is not a good candidate for selection if its expression is widespread across cells, or is expressed in a *random* set of cells, that is, a set of cells that do not have related transcriptomic profiles.

Let's suppose a scRNA-seq count matrix of $c$ cells by $g$ genes. Given a number $k$ of neighbors per cell (half the square root of the number of cells by default), *Triku* first creates a $k$-Nearest Neighbour ($K$NN) matrix using UMAP's `nearest_neighbors` function. To reduce computation times, the neighbour graph is calculated after applying PCA to the count matrix (30 components by default). The $k$NN matrix contains, for each cell in the matrix, the indices of its $k$ most similar cells.

Once the $k$NN matrix is calculated, the following procedure is applied for each gene. For each cell with positive read counts (read counts $> 0$), the number of counts of that cell and the number of counts of neighbour cells are summed. For each gene, this summarizes a distribution of the counts in the $k$NN cells. This is the $k$NN count distribution.

Next, the null distribution for each gene is calculated by choosing $k$ cells at random, and summing the counts together. To implement the random choices, we apply the convolution between read distributions. If $f$ is the density function of the distribution of reads in a gene, and $f'$ is the density function of the distribution of reads after removing the cells with no counts ($P(X = 0) = 0$), then the density function of $k$ random cells, one from $f'$ and $k$ from $f$ is $f'_k = f' * f * \cdots * f$, where $*$ is the convolution of functions, defined discretely as

$$(f * g)(X = i) = \sum_{m=-\infty}^{\infty} f(X = m)g(X = i - m)$$

.

Convolution assumes that cells are chosen with replacement, i.e. the same cell can be chosen more than once to calculate the distribution. Therefore, the distribution derived from the convolution is an approximation of the true null distribution, which considers no cell replacement. With single cell datasets, where the number of cells is high ($> 100$) and the value of $k$ is relatively small, the convolution creates an almost indistinguishable distribution compared to the null distribution.

##### 2.1.2 Comparing $k$NN count and null distributions by means of Wasserstein distance

Once the $k$NN and the null distributions are computed for each gene, the next step is to compare them. There are several functions to evaluate the difference between two distributions: distribution absolute difference, Kullback-Leiber divergence, Jensen-Shannon divergence, etc. The Wasserstein distance, coined in 1970 by R.L. Dobroshin after Leonid Vaseršteĭn introduced it in 1969; is also known as Earth Mover Distance, which answers the question of the minimum effort required to transform one distribution into the other, as an analogy of moving one pile of earth between two places.

Wasserstein distance has been found to perform best against other distribution comparison distances in tasks like image classification [4, 5], contour matching [6], or document classification in natural language processing [7]. The Wasserstein distance has also been applied as a loss metric in Generative Adversarial Networks (GANs) [8], improving over the standard metric, the Jensen-Shannon divergence, by avoiding model collapse during training. Recently, the Wasserstein distance has also been used in single-cell RNAseq analysis [9].

Wasserstein distance is scale-dependent, which poses a problem because genes are expressed throughout different orders of magnitude, and genes with highest expression profiles will show the biggest distances, regardless of the intrinsic similarity of observed and null distributions. To make the Wasserstein distance scale-independent, it is divided by the standard deviation of the null distribution:

$$D := \frac{D}{\sigma_{null}}$$

##### 2.1.3 Median correction and selection of features

During visual inspection of different datasets, we observed that in certain datasets the Wasserstein distances tend to slightly increase with the mean expression of the genes. However, throughout the expression range we could detect many genes having low Wasserstein distances, and some of them having higher distances, for a similar interval of mean expression. To correct for that bias the median correction is applied, that is, the gene expression range–in logarithmic scale–is divided into $w$ intervals (100 by default), and the median distance value of the genes in that interval is substracted to each gene.

Once the median correction is applied, the features with highest distances are selected. If the number of features to select is not decided by the user, this number is automatically calculated by setting a cutoff point to the Wasserstein distance. The cutoff point is selected by the line method, a simplification of the *kneedle method* [1]. Genes are ranked by their distance and create a hockey-stick curve. After drawing a straight line connecting the two ends of the curve, the cutoff point of the curve is the furthest from the line.

When implementing the algorithm to calculate the exposed cutoff point, we added the parameter $S$, $0 \leq |S| < 1$, which adjusts the cutoff point to vary the number of selected genes. If $C$ is the highest distance between the straight line and the curve,the new distance is set to $C \cdot |S|$. If $S \geq 0$, the point highest on the rank such that its distance is greater than $C \cdot |S|$ is chosen, thus selecting fewer genes, and vice-versa.

###### 2.1.4 Input and output

If the input object is a *scanpy* `adata` object, then the selected features are returned as boolean *pandas* series in `.obs['highly_variable']`, as done with *scanpy*'s `scanpy.pp.highly_variable_genes`. Additional `adata` variables are filled parameters used and distance values. If the input object is not an `adata` object, a dictionary with the Wasserstein distances, a boolean array with selected features and other properties from the analysis is returned.

##### 2.2 Parameter robustness

*Triku* is designed to a large amount of parameters. Triku's 4 most influential parameters in FS are seed, number of PCA components ($p$), $k$ in $k$NN, and number of windows for median correction ($w$).

The seed affects PCA and $k$NN graph creation, because their output is conditioned by initial random seed. The number of PCA components affects the $k$NN graph. $k$ is crucial because it directly influences the observed and null distributions. Generally, lower $k$ values tend to show a *locality* of the dataset, and infer larger sets of communities. On the other hand, higher values of $k$, apart from increasing computation time –approximately $O(n^2)$–, provide a more *global* view of the dataset, that is, expression patterns in larger sets of cells are favoured against expression patters shared by fewer cells.

Lastly, $w$ can yield absurd scenarios for extreme values. For instance, if the number of windows equals the number of features, each feature will be subtracted to itself, and the median corrected features will be 0. Generally, a low ($< 25$) number of windows will give a ragged or zigzaggy look to the values of the median corrected features.

To test the robustness of $p$, $k$ and $w$ for each parameter the other two parameters will be fixed with standard values, the standard value of the parameter will be compared to the rest of values. For example, when evaluating the robustness of $k$, *triku* is run with $p = 30$, $w = 100$, varying $k \in \{5, 10, 20, 50, 100, 150, 200, 400\}$. Assuming that the number of cells is 10000, the default value is $\sqrt{10000} = 100$. Then, the overlap degree between $k = 100$ and the rest of $k$ values for the first 500, 1500 and 2500 features is computed. Then, the procedure is repeated for $k = 100$ and $k = 10$, and so on, until $k = 400$. Since *triku* is run with 5 different seeds, its effect is tested indirectly because comparison between $k = 100$ and $k = 100$ will not yield a full overlap.

### Supplementary figures

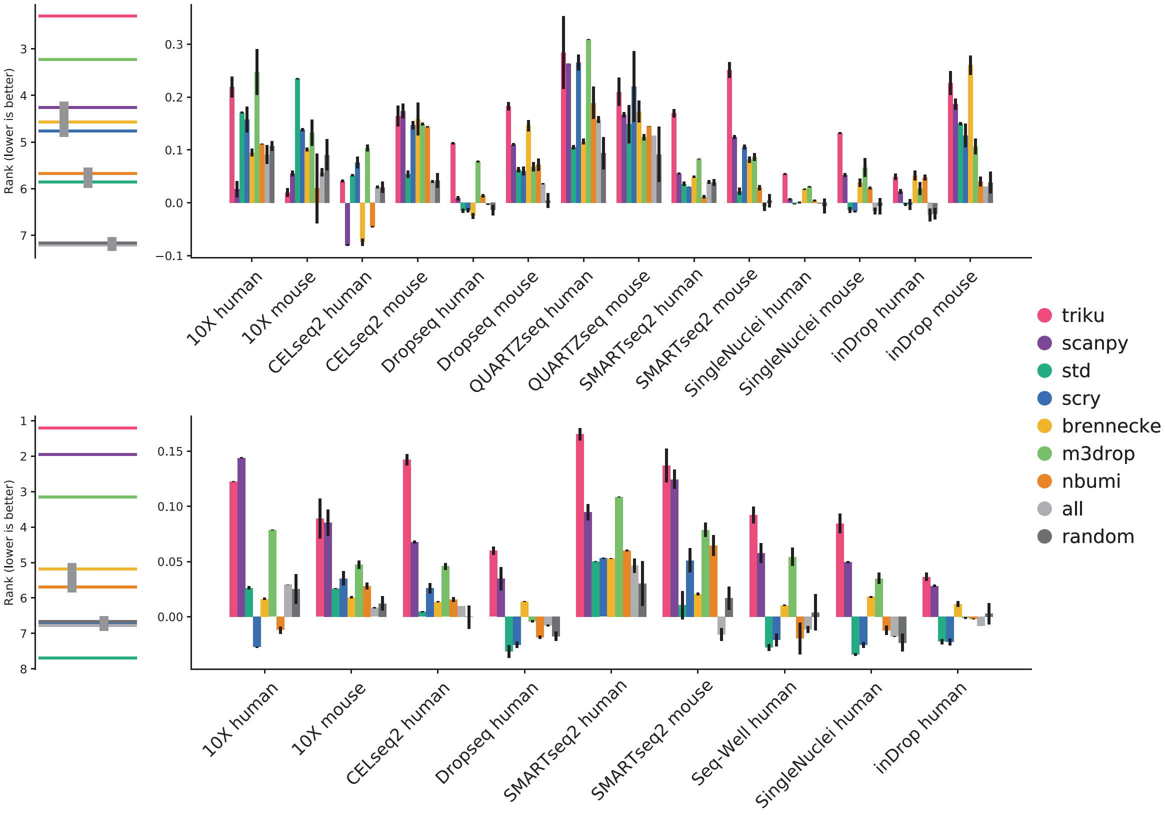


Figure S1: **Silhouette coefficients for leiden communities in Mereu and Ding datasets.** Barplots of Silhouette coefficient for Mereu (top) and Ding (bottom) datasets. Each barplot represents the mean of 5 seeds, and the vertical bar is the standard deviation. The plot on the left is a critical difference diagram, where each horizontal bar represents the mean rank for all datasets and all seeds. If two or more bars are linked by a vertical bar, the mean ranks for those FS methods are not significantly different (Quade test, $\alpha = 0.05$).

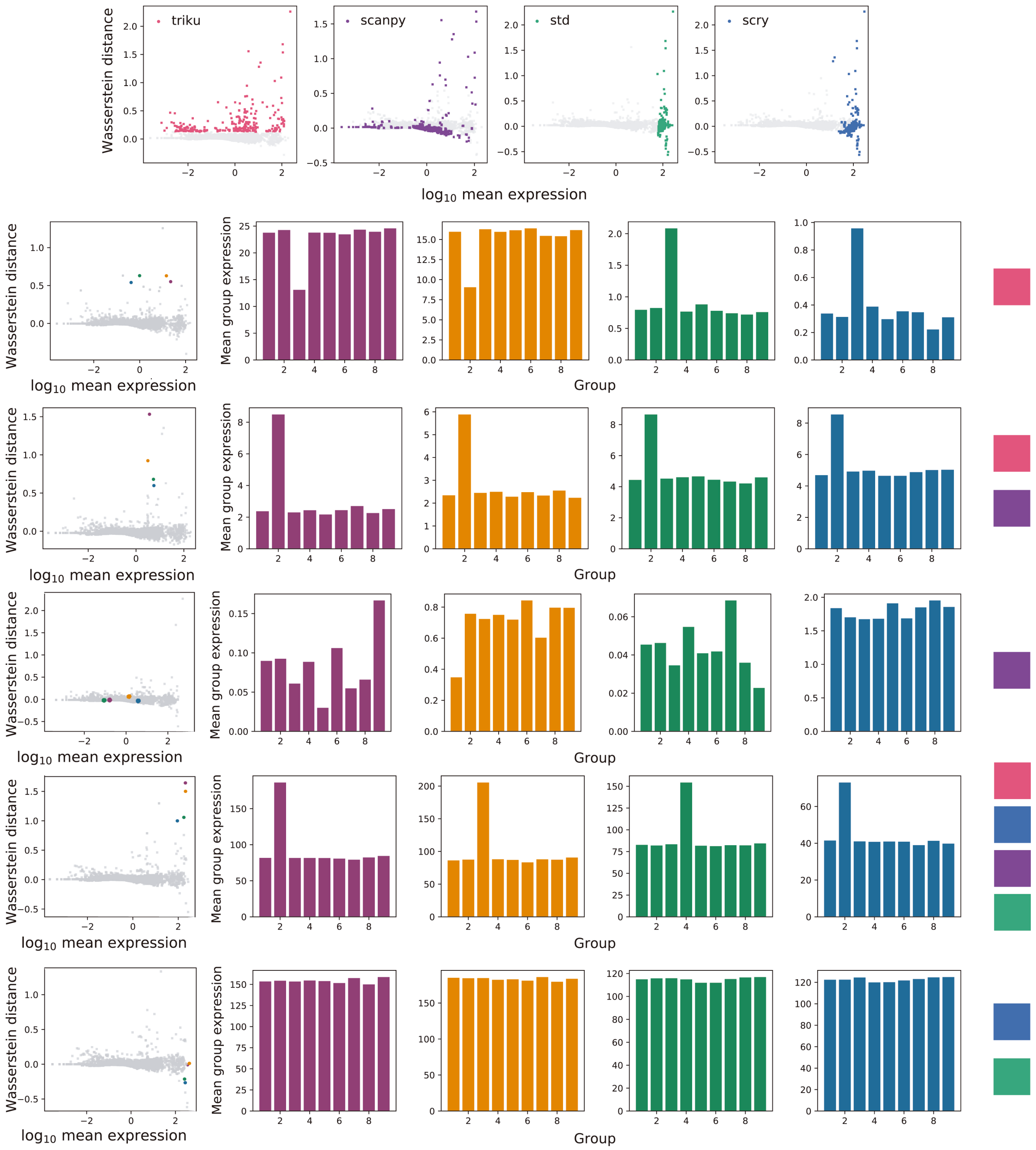


Figure S2: **Selected features for *triku*, *scanpy*, *std* and *scry* in artificial dataset with *de.prob* 0.01 and 250 genes.** The top row shows Wasserstein distance versus log mean expression scatter plots with the features selected by each FS method. The next rows show, for 4 genes, the mean expression per group of cells for each gene. The 4 features selected for each row are represented on the squares on the right: features selected only by *triku*, by *triku* and *scanpy*, by *scanpy*, by all FS methods, or by *std* and *scry*. We see that features selected by *triku*, with any other combination, have a group of cells where that gene is over or underexpressed, whereas features selected by other FS methods do not show groups with relevant over or underexpression.

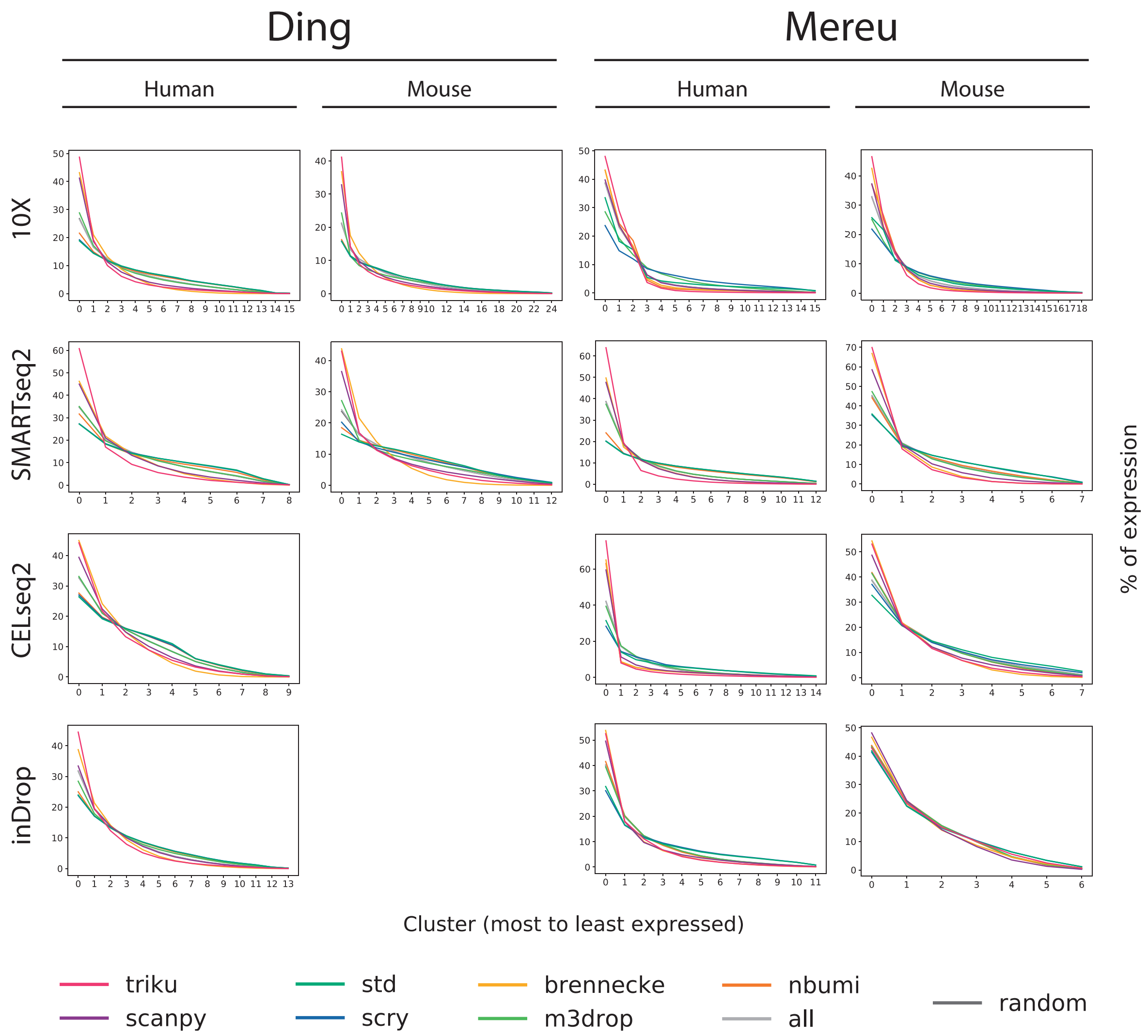


Figure S3: **Distribution of gene expression across clusters in *triku* is biased to fewer clusters.** For each of the datasets, and each gene, the expression of that gene was scaled to sum 1. Then, for each of the clusters obtained with leiden (resolution 1.2), the proportion of the whole expression is calculated, and the clusters are ranked, so that the cluster 0 has the highest proportion of expression compared to the rest of clusters. The lines in each plot represent the mean of the proportions for all selected genes for each FS method. For instance, in Ding's 10X human dataset, the most expressed cluster in features selected by *triku* express, on average, 50% of the expression of the gene, and the second most expressed one 20 %. Ding's CELseq2 and inDrop in mouse datasets do not exist, and are not shown in the figure.

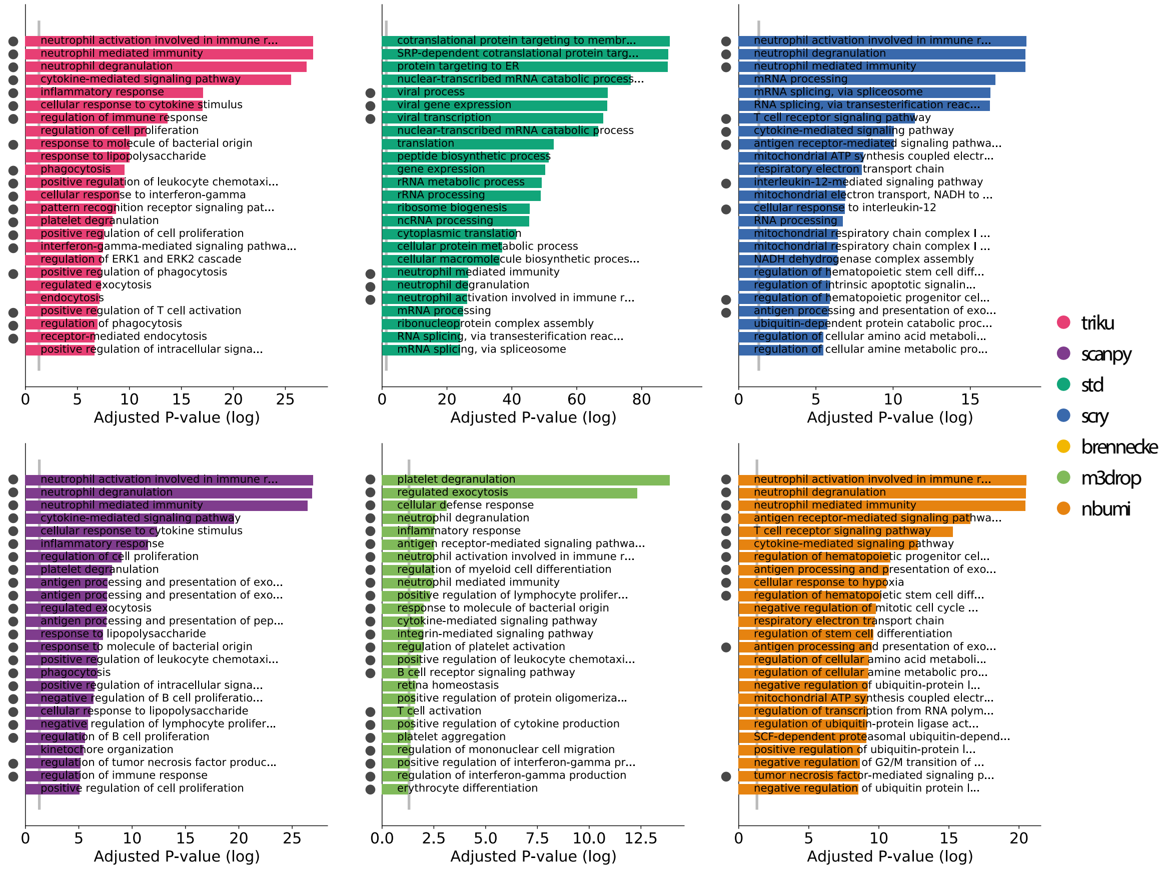


Figure S4: **Barplot of p-values of GOEA**. Each bin represents the number of features selected for each method, in Mereu et al. mouse 10X dataset. The represented value is the -log10 adjusted $p$-value for the best 25 ontologies. On the bottom, the bar plot shows the names of the ontology terms for the case with the best 1000 features. In immune datasets, gray dots at the left of each term represent that that term is directly-related to an immune process. Non-dotted terms refer to more general processes that may or may not be related to immune processes.

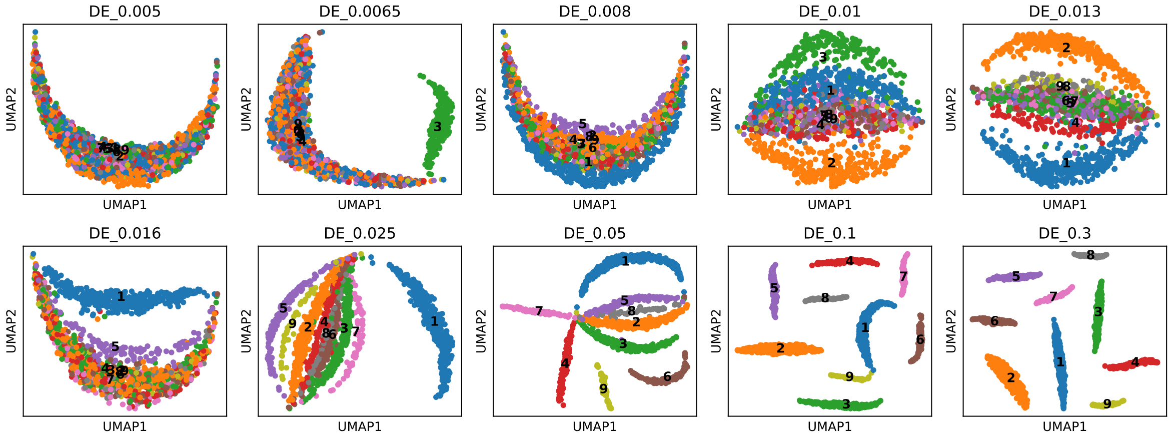


Figure S5: **Effect of scatter de.prob parameter on dimensionality reduction.** UMAPs of scatter datasets with different *de.prob* parameter values. UMAP and community detection were done without feature selection. Datasets with *de.prob* higher than 0.05 are completely resolved in UMAP, whereas lower values make scatter groups be less distinguishable.

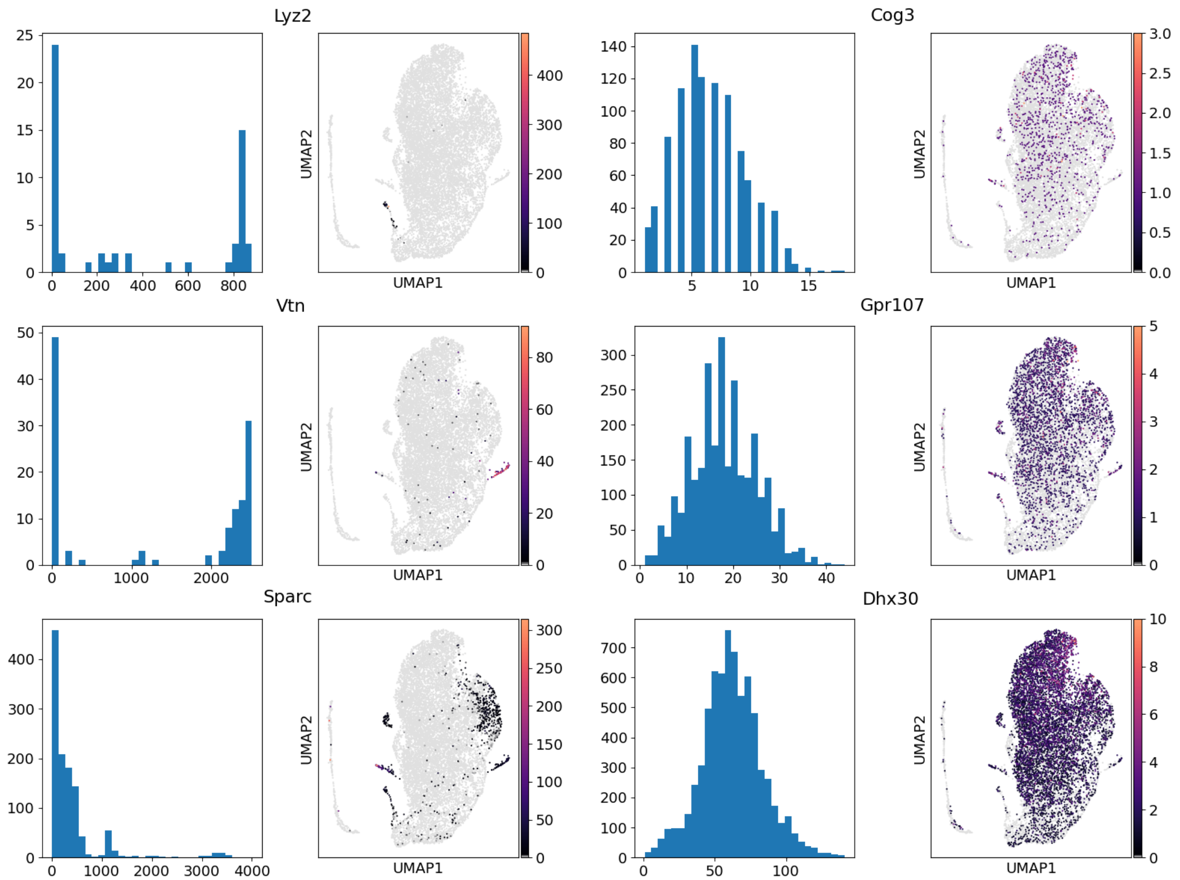


Figure S6: **Effect of proportion of zeros in gene expression patterns.** For each gene, the plot on the left represents the $k$NN count distribution of that gene, whereas the plot on the right is the UMAP DR representation with the $k$NN counts for each cell. The genes on the left (Lyz2, Vtn, Sparc) have a higher percentage of zeros and are, therefore, expressed in a subset of cells, whereas the genes on the right (Cog3, Gpr107, Dhx30) are more thoroughly expressed, and their $k$NN count distributions are not as heavy-tailed as for the genes on the left.

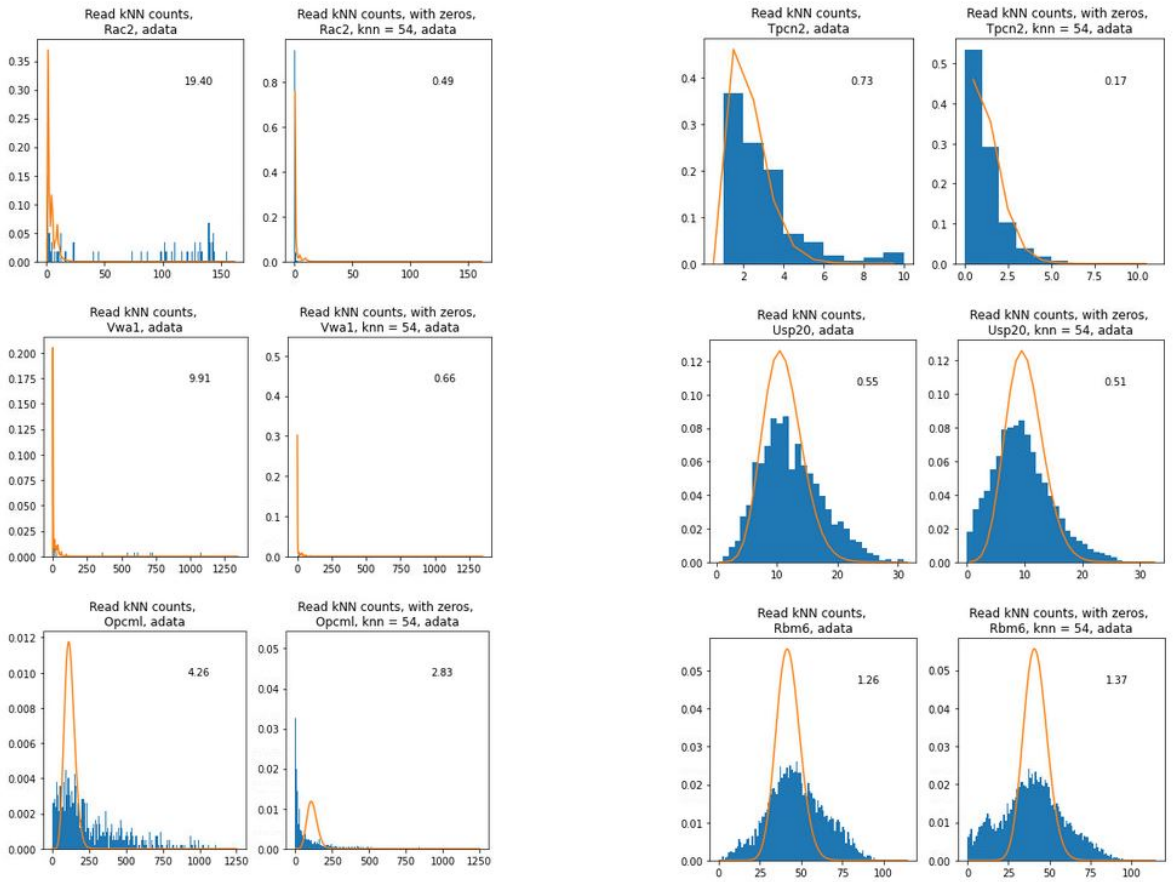


Figure S7: **Comparison of convolution and read distribution in $k$NNs.** The genes on the left (Rac2, Vwa1, and Opcml) are candidates to be selected by *triku*, whereas the genes on the right (Tpcn2, Usp20, Rbm6) are not candidates for selection. In each 6-block graph, the column on the left represents the counts –blue histogram– and convolution –orange line– of cells with positive expression, and their $k$NN; whereas the column on the right represents the counts and convolution of all cells, and their respective $k$NN. The number within each plot represents the Wasserstein distance between the convolution-based distribution and the $k$NN count distribution. The dataset used for this visualization is 10X neuron dataset, preprocessed in a similar fashion as the set of benchmarking datasets.

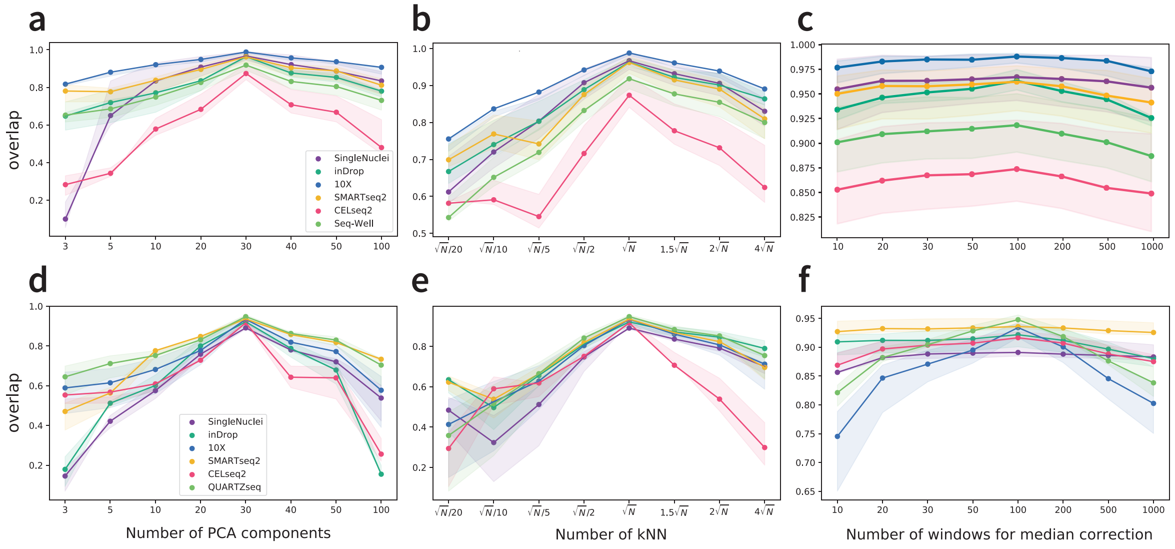


Figure S8: **Robustness of *triku* parameters.** Overlap values for the number of PCA components, $k$ and number of windows for benchmarking datasets from Ding et al. (a, b, c) and Mereu et al. (d, e, f). Lines in blue represent the overlap between the first 1000 features, and the extremes of the shaded regions represent the overlap between the first 500 (top) and 2500 (bottom) features.
